# LukProt: A database of eukaryotic predicted proteins designed for investigations of animal origins

**DOI:** 10.1101/2024.01.30.577650

**Authors:** Łukasz F. Sobala

## Abstract

The origins and early evolution of animals is a subject with many outstanding questions. One problem faced by researchers trying to answer them is the absence of a comprehensive database with sequences from non-bilaterians. Publicly available data is plentiful but scattered and often not associated with proper metadata. A new database presented in this paper, LukProt, is an attempt at solving this issue. The database contains protein sequences obtained mostly from genomic, transcriptomic and metagenomic studies and is an extension of EukProt (Richter et al., 2022, *Peer Community Journal*, **2**, e56). LukProt adopts the EukProt naming conventions and includes data from 216 additional animals. The database is associated with a taxonomic grouping (taxogroup) scheme suitable for studying early animal evolution. Minor updates to the database will contain species additions or metadata corrections and major updates will synchronize LukProt to each new version of EukProt and releases are permanently stored on Zenodo. A BLAST server to search the database is available at https://lukprot.hirszfeld.pl/. Users are invited to participate in maintaining and correcting LukProt. As it can be searched without downloading locally, the database can be a convenient resource not only for evolutionary biologists, but for the broader scientific community as well.

**Graphical abstract:** 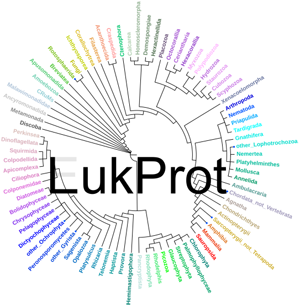

**Significance statement:** LukProt is a sequence database aiming to accelerate the research on the evolution of animals by cutting the time-consuming step of assembling sequences from disparate sources. Non-bilaterians are currently not well covered by general purpose databases, despite plentiful, public sequencing data. These data were integrated into a consistently curated database, presented here. It can be downloaded and used locally or used via a public BLAST search server. A clear taxonomic framework is also introduced, as well as scripts to aid local data analyses. LukProt will be publicly available on Zenodo, kept up to date and synchronized with each new version of its parent database, EukProt.

## Introduction

Animals (Metazoa) are a diverse group of organisms whose deep evolutionary history has been a subject of a long debate (Ruiz-Trillo et al., 2023). Many early branching lineages of animals are extinct, for example much of the Ediacaran fauna is not similar to any living animals. Animal tissues were not often amenable to fossilization pre-Ediacaran (Murdock & Donoghue, 2011). Recent advancements in phylogenomics helped resolve fundamental questions in early history of Metazoa, for example the identity of the earliest extant animal branch (Dunn et al., 2008; Schultz et al., 2023) and the timing of the two whole genome duplications which occurred in the ancestors of all living jawed vertebrates (Simakov et al., 2022; Marlétaz et al., 2024). These developments were enabled by more advanced bioinformatics tools, broader access to processing power and broader availability of sequencing data.

As animals are members of Eukaryota, knowledge about eukaryotic evolution can provide context for evolution of animals. In the last decade, the topology of the eukarotic tree of life has been undergoing revisions as major as the animal tree. The discovery and characterization of Asgard archaea (Zaremba-Niedzwiedzka et al., 2017; Eme et al., 2023) helped place the trunk of the tree. Due to the recent sequencing boom, new supergroups of protists are still being defined (Tikhonenkov et al., 2022). A comprehensive database which collects predicted proteomes of eukaryotes, EukProt, plays a role in these developments (Richter et al., 2022). The broad scope and careful curation of EukProt makes it useful for many applications, but the database compromises on the number of included species from mostly multicellular clades (plants, fungi or animals).

Investigating animal evolution requires broad taxon sampling of early diverging animal clades, which EukProt, NCBI or UniProt databases do not provide. One attempt at a wide animal taxon sampling, the Animal Proteome Database – AniProtDB (Barreira et al., 2021), includes only 100 species. Another large database of non-vertebrate animal sequences focuses on protostomes (Fernández et al., 2022). A more comprehensive database, which includes data from early diverging lineages, could serve as a tool for researchers to study the origins and evolution of uniquely animal characters, such as obligatory multicellularity or neurons.

LukProt, created to fill this gap, is a synthesis of the latest available EukProt, AniProtDB and many additional collected sequences. The database focuses on holozoans (a clade of opisthokonts sister to Nucletmycea) and animal species from the clades: Ctenophora, Porifera, Placozoa and Cnidaria. Bilaterian sampling is also increased. Conventions established by EukProt are followed: each dataset (proteome) has a unique identifier and each protein has a persistent ID composed of the dataset identifier, the species/strain name and protein number. A detailed characterization of LukProt, as well as EukProt data or metadata modifications made for LukProt, is presented below.

## Results and Discussion

### Purpose of LukProt

The purpose of the database release is to create a resource which one could consult on whether a given domain/protein/protein family exists within a clade of eukaryotes, with a focus on animals and their closest relatives: choanoflagellates, filozoans, ichthyosporeans and corallochytreans. To this end, individual dataset quality is of lower importance. No decontamination, or contamination-based dataset exclusion was performed. It is assumed that phylogeny reconstructions will be the guide when drawing conclusions from database searches. Phylogeny-guided analysis of LukProt, using a suitable sister database, can also be the basis for studies of lateral gene transfers. LukProt is documented with less experienced users in mind.

### Characterization of the LukProt database

The current version of the database (v1.5.1.rev1) contains 1281 datasets: 990 from EukProt, 41 from AniProtDB and 250 newly added. These newly added datasets were collected from various studies and publicly available repositories, or generated *in silico* from public data. Details of each dataset generation can be found in the metadata. Overall database statistics are given in **Supplementary Table 1**. Coverage of the early branching animal clades, especially ctenophores, sponges and cnidarians, is improved in comparison to EukProt. For example, 26% of the 23 currently known, potential placozoan species (Tessler et al., 2022) is included (6 datasets).

#### A consensus taxonomy

The database incorporates a consensus taxonomy of eukaryotes, largely based on UniEuk (Berney et al., 2017). The clades are explained in Supplementary Information and the full tree can be found in Figure 1. Each internal node was assigned a unique name, following UniEuk or recent literature where possible. Unnamed clades were assigned provisional names (Figure 1). The LukProt taxonomic framework will be adjusted to UniEuk updates and future data. It encompasses 81 groups of organisms, mostly monophyletic. The metadata also contains 59 groupings based on the consensus taxonomy, selected to aid targeted studies (e.g. Holozoa excluding Metazoa).

**Fig. 1.**
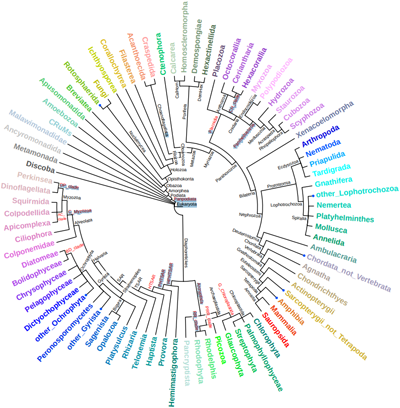
The eukaryotic tree of life used in LukProt. Each group is assigned a unique color which makes visual interpretation of phylogenies easier. Groups with blue dots are paraphyletic. Internal node names written in red were invented for the database.

#### Release schedule and data traceability

LukProt is available as an archive containing FASTA files placed inside a directory tree which recapitulates Figure 1. Additionally, archives with BLAST+ v5 database files (one per species, one per taxogroup and one full database) are available to download. The associated Zenodo repository is designed to be easy to use and contain files ready to be implemented in local workflows with minimal user modification required. These files include: the coloring scheme, the changelog, the metadata spreadsheet and helper scripts.

The latest database version is 1.5.1.rev1, based on EukProt v3. Major LukProt updates will trace each new release of EukProt, minor updates will be released as needed. The version will be incremented only if the underlying sequence files are changed; metadata changes will be marked “revisions”. All public versions of the database are archived in the Zenodo repository with a unique DOI.

Considerable effort was made to apply the traceability guidelines set by EukProt to the metadata included with LukProt. If available, the source URL and the article DOI is provided. As an improvement over EukProt, metadata also include NCBI taxids.

#### ‘LukProt Comparative Set’ (LCS)

By analogy to ‘The Comparative Set’ of EukProt – a reduced set of species of high sequencing quality and phylogenetic importance – the LCS is proposed. The set consists of 233 datasets, 18% of the total number, and is available to search using the BLAST server.

#### BLAST server and convenience scripts

A BLAST+ (Camacho et al., 2009) server for LukProt, available at https://lukprot.hirszfeld.pl/, utilizes SequenceServer 3.1.2 (Priyam et al., 2019) and NCBI BLAST+ 2.16.0+ and incorporates the tree structure from Figure 1. The accompanying scripts are documented with usage examples and can be used to: recolor tree tips, extract sequences from preselected groups of organisms (including the LCS) or extract LukProt sequences from the full BLAST database or using a list of identifiers.

### LukProt data sample: top single BUSCOs analysis

Figure 2A presents an example of a phylogeny reconstructed from 20 most often found single eukaryotic BUSCOs in the database. No post-tree reconstruction sequence removal was performed for a fairly unfiltered view of data found in LukProt for the selected clade, Filozoa.

**Fig. 2.**
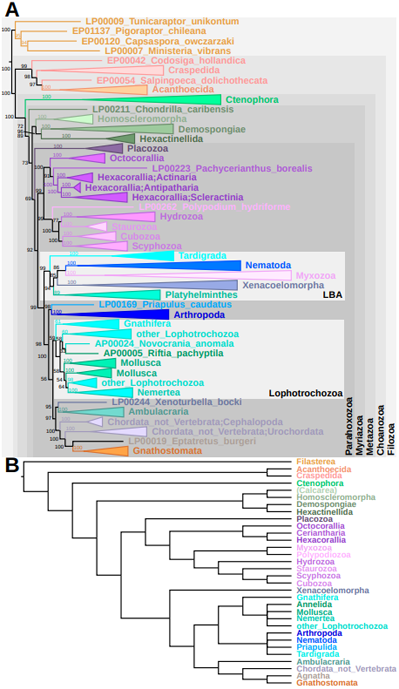
(**A**) Consensus IQ-TREE phylogeny reconstruction of the 20 most prevalent LukProt BUSCOs in Filozoans. UFBoot2 branch support values are indicated. LBA – long branch attraction artifact. (**B**) Reference, colored cladogram of the group Filozoa. Calcarea is shown in brackets because of its absence from the main phylogeny.

The phylogeny in Figure 2A mostly agrees with the current understanding of filozoan/early animal evolution (Figure 2B). Ctenophores were recovered as the earliest branching animal clade with full support. Most sponge sequences formed a single clade, as well as Cnidaria and Bilateria. However, the long the branch attraction phenomenon (Susko & Roger, 2021) caused formation of a branch with animals from clades known to have highly divergent sequences.

The alignment used here contained 299 species and 5229 parsimony informative sites. If sequences are short, a search of LukProt may return too many for reliable maximum likelihood inference (trees may be overparametrized). In such cases, sequence similarity networks (SSNpipe – https://github.com/ahvdk/SSNpipe), cd-hit or MMSEQS2 (Mirdita et al., 2019) clustering can be used to reduce their number.

### Limitations of the LukProt database and future plans

The limitations of the parent database (EukProt) apply to LukProt as well. One further limitation results from a decision not to predict proteins shorter than 100 amino acids (50 in EukProt). The LCS is a beta version and the community is invited to suggest changes. Any missing NCBI taxids will be added in due course.

## Materials and Methods

### Dataset naming and species selection

In most cases, species and strain names from EukProt were conserved. To aid further analyses, extremely long species/strain names were shortened so that no species+strain name is longer than 32 characters (including underscores). In these cases, only the organism name was truncated – the EukProt dataset identifier and the protein ID were not changed. The name mapping file is available in the Zenodo repository. Sequence names from source files are included after each LukProt ID.

Newly downloaded datasets were assigned names after reading the associated publications, if any referred to the name. Otherwise, names were adapted from the websites from which the datasets were downloaded. As in EukProt IDs, spaces are replaced with underscores in individual FASTA files and additionally periods were removed from the names. To broaden compatibility and reduce file size, the following additional processing steps were performed on the datasets from EukProt: removing asterisk (*) characters and line breaks from sequences.

### Data processing – LukProt

The data processing steps and software versions are detailed in the associated metadata file. Sequence manipulation was done using SeqKit (Shen et al., 2016) and GNU coreutils. Unassembled transcriptomes were assembled using Trinity v2.12 to v2.15.1 (Grabherr et al., 2011; Haas et al., 2013) or TransAbyss v2.0.1 (Robertson et al., 2010). TransAbyss was used 1 case where Trinity returned an error. Proteins were predicted using TransDecoder.LongOrfs (Haas, BJ. https://github.com/TransDecoder/TransDecoder, versions v5.5.0/v5.7.0). Predicted peptides were clustered (in most cases) using cd-hit v4.8.1 (Fu et al., 2012). Strains were merged by running cd-hit on the concatenated FASTA files. Gffread v0.12.7 (Pertea & Pertea, 2020) was used to extract proteins from annotated genomes. NCBI taxonomy IDs were found manually or with the R package taxonomizr (Sherrill-Mix, S. https://github.com/sherrillmix/taxonomizr v0.10.6). BUSCO 5.7.1 (Manni et al., 2021) was used with dataset “eukaryota_odb10” (255 BUSCOs) created on 2024-01-08. BUSCO utilized hmmsearch version 3.1 (http://hmmer.org/). The OMArk (Nevers et al., 2024) version was 2.0.3 and the OMAmer version was 0.3.0, used with database LUCA.h5 (Nov 2022). The NCBI taxdb used with OMArk was downloaded on 2024-05-12. BUSCO, OMAmer and OMArk were run with default parameters and the species NCBI taxid was specified for OMArk where available.

### Data processing – 20 BUSCOs example

First, 20 most common single BUSCOs in the LukProt database were selected. Sequences of these 20 BUSCOs were collected into separate FASTA files for each dataset. Species which possessed fewer than 10 of these BUSCOs were filtered out. BUSCO sequences from the remaining species were then concatenated, merged into a single multi-FASTA file and aligned using FAMSA v2.2.2-(Deorowicz et al., 2016). The alignment was trimmed using trimAl v1.4.rev15 (Capella-Gutierrez et al., 2009) (gap threshold 0.5). Phylogenies were inferred using IQ-TREE 2.3.3. IQ-TREE branch support values were calculated using SH-aLRT (Shimodaira–Hasegawa-like approximate likelihood ratio test, 5000 replicates) (Guindon et al., 2010), aBayes (approximate Bayes) (Anisimova et al., 2011) and UFBoot2 (Ultrafast bootstrap 2, 5000 replicates) (Hoang et al., 2018). IQ-TREE model was selected using ModelFinder (Kalyaanamoorthy et al., 2017). FigTree 1.4.4 (http://tree.bio.ed.ac.uk/software/figtree/) was used for phylogeny visualization and editing.

## Acknowledgements

I would like to thank the maintainers of the EukProt database. In addition, Marcin Czerwiński, Marta Álvarez, Elena Casacuberta, Michelle Leger, Koryu Kin and Iñaki Ruiz-Trillo, are thanked for helpful discussions and permission to release the *Corallochytrium limacisporum* (strain India) proteome. I further thank Michelle Leger and Eduard Ocaña-Pallarès for suggestions about the taxonomic framework.

This work was supported by National Science Centre of Poland (NCN) as part of the grant SONATINA no. 2020/36/C/NZ8/00081 entitled “The role of glycosylation in the emergence of animal multicellularity”.

## Data availability

The data underlying this article (the LukProt database and associated files) are available on Zenodo: https://doi.org/10.5281/zenodo.7089120 (Sobala, 2022) and https://doi.org/10.5281/zenodo.11324807 (Sobala, 2024).

