## Supplementary Information for "LukProt: A database of eukaryotic predicted proteins designed for investigations of animal origins"

### Supplementary Tables

Supplementary Table 1 – Comparison of the number of taxa in selected clades of LukProt and other databases.

| Taxogroup | EukProt v3 – included in the stock database | AniProtDB within LukProt | LukProt v1.5.1.rev1 added (+)/excluded (-) | LukProt v1.5.1.rev1 total |
|---|---|---|---|---|
| Eukaryota | 993 | 41 | +250/-3 | 1281 |
| Diaphoretickes | 632 | 0 | +28/-0 | 660 |
| Holozoa (excluding Metazoa) | 105 (40) | 41 (0) | +221/-2 (+5/-1) | 365 (44) |
| Metazoa (excluding Bilateria) | 65 (14) | 41 (3) | +216/-1 (+162/-0) | 321 (179) |
| Ctenophora | 2 | 1 | +35/-0 | 38 |
| Porifera | 5 | 1 | +41/-0 | 47 |
| Placozoa | 2 | 0 | +4/-0 | 6 |
| Cnidaria | 5 | 1 | +82/-0 | 88 |
| Bilateria | 51 | 38 | +54/-1 | 142 |

### Supplementary Materials and Methods

Command line options used with bioinformatics software, where different from defaults:

CD-HIT: cd-hit -s 0.5 -g 1 -d 0 -T 6 -M 16000 -c 0.99 -i input.fa -o output.fa

Gffread: gffread -g genome.fasta annotation.gff -y output.fa -C -J –no-pseudo

FAMSA: famsa -refine-mode on

### Clade naming conventions

The taxonomy is based on multiple sources. Deep cladistic relationships were inferred from: (Cavalier-Smith, 2013; Ruggiero et al., 2015; Heiss et al., 2018; Lax et al., 2018; Strassert et al., 2021; Tikhonenkov et al., 2022). Selected clades are listed below in a tree-like structure and sourced where appropriate. Items with an asterisk are convenience names invented for the purpose of this database. Part of the invented names will be submitted to UniEuk for consideration by the community.

Eukaryota: Ancyromonadida + Diaphoretickes + Discoba + Metamonada + Panpodiata*
 Diaphoretickes: Arcryptista* + HPHTSAR*
  Arcryptista*: Archaeplastida + Pancryptista
   Archaeplastida: G_Chloroplastida* + PRR_clade*
    G_Chloroplastida*: Glaucophyta + Chloroplastida

PRR_clade*: Picozoa + RR_clade*
RR_clade*: Rhodelphis + Rhodophyta
Pancryptista (Yazaki et al., 2022): Cryptista + *Microheliella maris*
HPHTSAR* (Tikhonenkov et al., 2022): Hemimastigophora + PHTSAR*
PHTSAR* (Tikhonenkov et al., 2022): Provora + HTSAR*
HTSAR* (Tikhonenkov et al., 2022): Haptista + TSAR
TSAR: Telonemia + SAR
SAR: Halvaria + Rhizaria
Halvaria (Ruggiero et al., 2015): Alveolata + Stramenopiles
Alveolata: Colponemidae + C_Myzozoa*
C_Myzozoa*: Ciliophora + Myzozoa
Myzozoa: AC_clade* + DP_clade*
AC_clade*: Apicomplexa + Colpodellida + Squirmida
DP_clade*: Dinoflagellata + Perkinsea
Stramenopiles: Gyrista + Bigyra + Platysulcus
Gyrista: Ochrophyta + Peronosporomycetes + other_Gyrista
Ochrophyta: BD_clade* + Chrysophyceae + Pelagophyceae + Dictyophyceae + other_Ochrophyta
BD_clade*: Bolidophyceae + Diatomeae
Panpodiata* (Tikhonenkov et al., 2022): Malawimonadidae + Podiata
Podiata: CRuMs + Amorphea
Amorphea: Obazoa + Amoebozoa
Obazoa: Opisthokonta + Breviatea + Apusomonadida
Opisthokonta: Holozoa + Nucletmycea
Holozoa: Ichthyosporea + Corallochytrea + Filozoa
Filozoa: Filasterea + Choanozoa
Choanozoa: Choanoflagellata + Metazoa
Metazoa (Schultz et al., 2023): Ctenophora + Myriazoa
Myriazoa (Schultz et al., 2023): Porifera + Parahoxozoa
Parahoxozoa (Schultz et al., 2023): Bilateria + Placnidia
Placnidia* (Laumer et al., 2018; Schultz et al., 2023): Placozoa + Cnidaria
Cnidaria: Anthozoa + Panmedusozoa*
Anthozoa: Octocorallia + CH_clade*
CH_clade*: Ceriantharia + Hexacorallia
Panmedusozoa*: Endocnidozoa + Medusozoa
Endocnidozoa (Kayal et al., 2018; Koch et al., 2021; Xiao et al., 2022): Myxozoa + Polypodiozoa
Medusozoa (Kayal et al., 2018; Klompen et al., 2021): Hydrozoa + Acraspeda
Acraspeda (Kayal et al., 2018, p. 201): Staurozoa + Rhopaliophora
Rhopaliophora (Kayal et al., 2018): Cubozoa + Scyphozoa
Bilateria: Nephrozoa + Xenacoelomorpha
Nephrozoa (Cannon et al., 2016): Protostomia + Deuterostomia
Protostomia: Ecdysozoa + Lophotrochozoa
Ecdysozoa (Marlétaz et al., 2019): Arthropoda + Nematoda + Priapulida + Tardigrada
Lophotrochozoa (Marlétaz et al., 2019): Gnathifera + Spiralia

Spiralia (Marlétaz et al., 2019): Annelida + Mollusca + Nemertea + Platyhelminthes + other_Lophotrochozoa
